# Microplastics cross the murine intestine and induce inflammatory cell death after phagocytosis by human monocytes and neutrophils

**DOI:** 10.1101/2025.03.10.642492

**Authors:** Giulio Giustarini, Tim L.P. Skrabanja, Annemijne E.T. van den Berg, Selma van Staveren, Tom Vos, Zhi Hui Zhou, Thomas Klaessens, Eva Mulder, Joëlle Klazen, Joost Smit, Raymond Pieters, Leo Koenderman, Nienke Vrisekoop

## Abstract

Microplastics are contaminating the environment but also our food and drinking products. In a crucial study, microplastics have been detected in circulating human blood, urging the investigation of the effects of microplastics on human health. Here we aimed to determine the distribution of microplastics after oral exposure in mice and their interactions with and effects on mouse and human innate immune cells. We established that both 1 and 10μm polystyrene (PS) particles penetrated the intestinal epithelium after oral administration and could be detected in blood and liver of mice after ten days of oral administration. Using intravital microscopy we captured the in vivo phagocytosis of 1μm PS by mouse neutrophils in the liver. Pristine PS were barely phagocytosed by primary human phagocytes, however, 1 and 10µm PS pre-incubated with plasma were readily phagocytosed by human neutrophils and monocytes. Plasma-coated 1 and 10μm PS both increased human neutrophil and monocyte cell death but only after phagocytosis. Importantly, neutrophil cell death occurred a few hours after phagocytosing a single coated 10µm PS while PS of 1µm needed to be administered at a ratio of 27 particles per cell to induce significant neutrophil cell death. Neutrophil cell death upon microplastic exposure was characterized by extracellular DNA, which together with other released DAMPs can potentially trigger inflammation. Our findings suggest that microplastics could negatively impact the immune system and human health.

## Introduction

Plastics have bestowed considerable advantages to humanity. The production of plastics has been steadily increasing for more than half a century. In 1950 the total plastic production was approximately 1.9 million tons, rising to an approximate 400 million tons in 2022 (PlasticsEurope AISBL). Vast and considerable amounts of plastics are released or abandoned in the environment. These plastics are either intentionally produced as small particles or gradually fragmented into smaller particles by the mechanical action of waves and currents, the uptake by birds and fish, chemical processes such as oxidation and hydrolysis, and exposure to UV light^3–5^. Thus, most plastics deposited in the environment persist as ‘microplastics’, a term coined by Professor Richard Thompson in 2004^6^. Microplastics have been detected in air, aquatic life, drinking water, and several foods and beverages^1,2 ,7–11^.

As highlighted by the WHO, not much is known on human exposure and the risk represented by microplastics for human health^12^. Some studies estimate that humans may ingest as much as 5 g of plastic per week^13,14^. However, estimates heavily depend on unknown assumptions and/or preliminary non-standardized data^15,16^.

Intestinal translocation to gut, liver and kidney of both 5 and 20µm polystyrene (PS) microplastic particles is reported in mice after daily oral administration of 0.1mg^14^. In line, daily administration of 0.2mg 50nm or 1µm PS for 24 weeks via drinking water resulted in translocation in the small intestine and presence of PS in mesenteric lymph nodes, spleen and liver^17^. In contrast, Stock et al. orally administered 1µm (25µg/kg), 4µm (1.6mg/kg) and 10µm (819 µg/kg) PS particles 3 times per week and only detected very few 1µm particles in the intestine and none in liver, spleen and kidney^18^. Regardless, microplastics have been detected in human blood with a mean concentration of 1.6 µg/mL blood^19^, demonstrating microplastics translocate over the intestinal and/or lung epithelial lining in vivo in humans. The size of the microplastic particles in human blood is, however, still unknown.

Our immune system defends our body against foreign particles, like pathogens. Immune cells can recognize patterns on pathogens, engulf/ phagocytose them and subsequently degrade these threats. Microplastics that enter our body via the gastro-intestinal tract might also be considered foreign, however, our immune system has not evolved to recognize patterns on microplastics. Limited studies have been performed with microplastics and professional phagocytes. Neutrophils were shown to phagocytose 1µm PS in mice after intravenous administration^20^. Furthermore, neutrophils phagocytosed 1µm PS in lung, liver and kidney after oral administration in mice^21^. In line, human neutrophils were shown to phagocytose 1µm PS in vitro. Phagocytosis led to neutrophil death in both mouse and human^21^. These preliminary data prompt more in-depth research into the distribution of microplastics after oral exposure and their interactions with and effects on mouse and human innate immune cells.

In this study, we assessed the fate of microplastics after oral exposure to 1 and 10µm PS particles. To this end, we investigated if 1 and 10µm PS particles were transferred across the epithelial intestinal barrier after oral exposure in mice. Additionally, we examined the interaction between PS particles and professional phagocytes which are part of the first line of defense of the immune system. We intravitally imaged phagocytosis of 1µm PS microplastics by neutrophils in the liver of mice. Furthermore, we monitored the kinetics of in vitro phagocytosis and cell survival of human neutrophils and monocytes after exposure to different concentrations of 1 and 10µm PS particles.

## Methods

### Microplastic PS particles, particle coating and pH-rodo labeling of particles

Standard non-fluorescent and fluorescent (Ex/Em 502 nm/518nm) PS microplastic particles of 1, 5, 10µm were purchased from Microparticles GmbH (Berlin, Germany).

Particles were incubated with plasma, serum, human serum albumin (HSA) or their heat-inactivated (HI) form (only serum and HSA) at 37°C for 2 h. After incubation 10µm particles were washed with incubation buffer and resuspended. Particles below 5µm diameter were diluted to reach a max concentration of 5% of plasma, serum or HSA during the assays. All the conditions were equilibrated for the content of serum, plasma or HSA depending on the experiment.

Particles were spun down at 10,000g and resuspended in 0.1M NaHCO3. An aliquot of desiccated pHRodo^TM^ Red (Invitrogen) was resuspended in DMSO according to manufacturer’s protocol. pHRodo^TM^ was incubated to a final concentration of 0.5mM for 1 hour in the dark at room temperature. The particles were washed with PBS and resuspended in incubation buffer to achieve a concentration of 2.5% W/V.

### Mouse experiments Mice

Male, 9 to 11-week old, C57BL/6J mice (The Jackson Laboratory, Charles River) were used for all experiments. They were allowed to acclimatize for 1 week in a 12-h light/dark cycle and maintained at mean temperature of 23 ± 2°C, 50–55% relative humidity. Drinking water and laboratory food pellets were provided ad libitum. In vivo studies were approved by the Ethics Committee for Animal Experiments of Utrecht University and complied with governmental and international guidelines on animal experimentation.

### Oral administration of PS microplastics

For 1-10 consecutive days mice received an intra-gastric gavage delivering a suspension of 4mg of either green or red fluorescent PS (1 and 10µm diameter) in 160 µl of water. Control mice received only the vehicle. Blood was collected from these mice 0.5, 2 and 24h after the first intra-gastric administration and 24h after the last gavage on day 10. After blood collection mice were sacrificed and gut and liver were excised and prepared for histology, snap frozen and stored at -80 °C till use.

### Flow cytometry

Two hundred µl of whole blood was spun at 300g for 10 min after which 50µl plasma was collected and further plasma aspirated. Red blood cells were lysed (Lysis buffer: 155mM NH_4_Cl, 10 mM KHCO_3_, 0.1mM EDTA) and the pelleted cells resuspended in 150µl of incubation buffer (20mM Hepes, 132mM NaCl, 6mM KCl, 1mM MgSO_4_, 1.2mM KH_2_PO_4_, supplemented with 1.0mM CaCl_2_, 5mM glucose, 0.5% [w/v] HSA). Red or green fluorescent particles were detected in both plasma and cells using a Canto/ Fortessa flow cytometer (BD).

### Histological immunofluorescent staining

Immunofluorescence was performed on frozen liver and intestinal tissues embedded in optimal cutting temperature compound. Eight μm thick frozen sections were mounted on polylysine-coated glass slides, dried overnight at room temperature and stored at –20°C until use. Briefly, tissue sections were allowed to dry for 2 hours before fixation with 4% formaldehyde in PBS. Intestinal samples were stained using Vector Red Alkaline Phosphatase substrate following procedures reported in manufacturer’s instructions (cat. N. SK-5100). Nuclei were identified by DAPI staining after permeabilization with PBS containing 0.5% Triton-X100 for 5 minutes.

### Intravital microscopy

For intravital microscopy, Lys-M-EGFP mice received either the same intragastric dose of red fluorescent PS as mentioned above for 2 consecutive days or were injected retro-orbitally with 50µl of saline containing 3x10^7^ red fluorescent 1µm PS. After administration, Lys-M-EGFP mice underwent anesthesia with isoflurane. Under anesthesia mice received a subcutaneous injection of saline before starting the surgical preparation of the liver for intravital microscopy with a LSM-880 2-photon microscope (Zeiss, Germany). Before imaging, mice received an injection of Alexa Fluor 647-conjugated bovine serum albumin (BSA) to allow visualization of blood vessels.

### Experiments with human samples

Ethical approval for obtaining healthy human volunteer blood was obtained by the institutional ethical review board of the University Medical Centre Utrecht (UMCU) under protocol number 07-125/C and all subjects provided written informed consent.

### Isolation of blood neutrophils and monocytes Neutrophils

Neutrophils were isolated from fresh whole blood as described previously by Overbeek et al. (2013). Neutrophil preparations consisted of 95–97% neutrophils as determined by an Abbot Cell-Dyn emerald.

Neutrophils were resuspended in incubation buffer (20mM Hepes, 132mM NaCl, 6mM KCl, 1mM MgSO_4_, 1.2mM KH_2_PO_4_, supplemented with 1.0mM CaCl_2_, 5mM glucose, 0.5% [w/v] HSA).

### Monocytes

Monocytes were isolated from peripheral blood mononuclear cells (PBMC) by adherence after isolation with Ficoll/Hystopaque 1.077 g/ml, as previously described ^22^. One million PBMCs were incubated directly in 96 well plates and let adhere for 2 hours at 37°C resulting in approximately 2x10^5^ monocytes per well. Non-adherent cells were removed by washing 3 times with incubation buffer after adherence ^23^.

Incucyte^®^ live cell analysis Phagocytosis

Neutrophils and monocytes (10^7^ cells/ml) were exposed to incubation buffer (without HSA) containing 10nM SNARF^®^-1 carboxylic acid, acetate, succinimidyl ester and placed on a rotating shaker for 5 min in the dark at room temperature (RT). Cell suspension was diluted 1:2 with pooled human serum to quench the staining (Sigma-Aldrich, MO, USA) and further diluted 10x with incubation buffer before centrifugation at 1500rpm for 5min at RT. Next, cells were resuspended in incubation buffer. Green fluorescent microparticles with and without coating were used and phagocytosis was analyzed with Incucyte^®^ analysis software. Briefly, SNARF^®^-1 fluorescence was used to create a mask to detect the area of the cells and phagocytosis was calculated as the area of merge between the fluorescence of the PS and the mask of the cells. Fluorescence was acquired in 4 different areas of each well.

### Cell viability

Neutrophils and monocytes were seeded in 96 well plates at a concentration of 2x10^5^ cells in 200µl in incubation buffer containing Annexin V (AnnV) (BioLegend, Uithoorn, The Netherlands) and propidium iodide (PI) (Sigma-Aldrich, St.Louis, Missouri). Non-fluorescent particles with and without coating were used at different indicated concentrations and cell:bead ratios. Fluorescence of AnnV and PI were detected every hour for 72h using the Incucyte®.

### Flow cytometry cell sorting

Isolated neutrophils (1 x 10^6^ cells/ml) were exposed to coated 10µm fluorescent particles for 1 hour in presence of f-MLF (10^-8^M) (Sigma-Aldrich, MO, USA). At this time, cells were prepared for cell sorting using an Aria III (BD Biosciences, NJ, USA). Fluorescent positive and fluorescent negative cells were seeded in different wells of 384 well plates (300 cells per well) containing incubation buffer to measure cell viability. In the wells containing fluorescent negative cells, an equal number of 10µm fluorescent particles were also seeded. Viability was measured in Incucyte® as mentioned above.

### Cell death characterization

For the cell death characterization, neutrophils were kept in Hepes3+ medium containing 4μM Hoechst 33342 (Merck Life Science N.V., 14533, Germany), 1μM Sytox blue (Fischer Scientific, S34857, United States) and AnnexinV-FITC (BioLegend, 640906, United States) in a black/clear polystyrene flat bottom imaging 96-well plate for imaging. Neutrophils were separately challenged with 0.05% (w/v) Triton-X100 (Merck Life Science N.V., T8787, Germany), 10μM Staurosporin (protein kinase inhibitor; Sigma- Aldrich, S4400, United States), 10μM Nigericin (NLRP3 inflammasome inducer; Merck Life Science N.V., N7143, Germany), 25ng/ml phorbol 12-myristate 13-acetate (PMA) (protein kinase C stimulator; Sigma-Aldrich, P8139, United States), or 137.44μg/ml serum-coated 10µm PS (1:1 neutrophil to PS ratio). Untreated neutrophils acted as a control. The imaging plate was live imaged on a Leica Stellaris 5 microscope at 20X magnification and 2.0 zoom at 37°C and 5% CO_2_. Every 10 minutes, 2 sets of images of each well were taken for 10 hours. Acquired images were processed with Fiji software. All experiments were done in duplo and the experiment was performed with three independent donors.

## Results

### Orally administered 1 and 10 µm fluorescent PS translocate to blood and liver in mice

Previous studies show that oral administration of PS particles leads to their accumulation in several organs like the liver, kidney and gut^24^. First, we aimed to confirm that 1 and 10µm fluorescent PS particles penetrate the intestinal epithelium and translocate to the blood. To this end, we orally administered these particles (115 mg/kg/day, ±4 mg/day to each mouse) on a daily basis for 10 consecutive days. This daily dose is about 10 times higher than the estimated exposure in humans^13^ as is common practice in toxicology^25^.

We first explored the translocation of orally administered particles in the small intestine by microscopy. The lumen of the small intestine contained several fluorescent particles of both 1 and 10µm, 2h after the first oral administration. Importantly, at this time point, it was possible to observe both 1 and 10µm green fluorescent polystyrene particles crossing the mucosa of the small intestine (Fig. 1).

**Figure 1.**
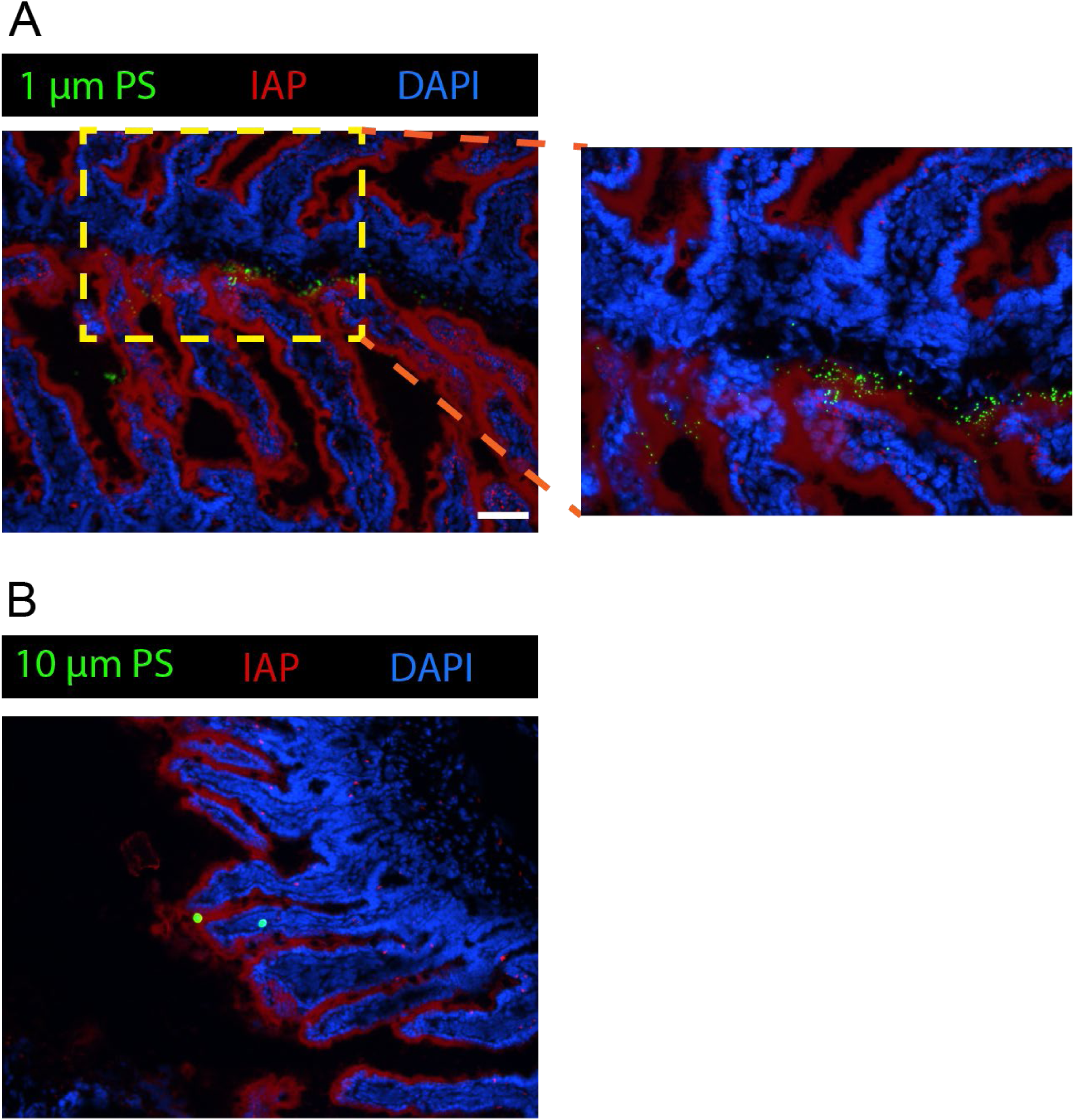
Intestinal translocation of 1µm and 10µm PS particles 2h after oral administration Intestinal section showing translocation of green 1µm (A) and 10 µm (B) PS particles over the intestinal lining stained by intestinal alkaline phosphatase (IAP) in red. Nuclei are stained blue using DAPI. Scale bar is 50 µm.

Blood was collected 0.5, 2 and 24h as well as 10 days after the first oral administration. Plasma (50µl) and the cellular fraction (from 200µl) were analyzed by flow cytometry. The 1µm fluorescent PS were detected in plasma as early as 30 minutes after the first oral administration as shown in Fig. 2A, although not significantly higher than the background measured in controls. Two hours after the first administration the 1µm particle count was significantly higher than the control. These particles were no longer detectable in plasma 24h after administration (Fig. 2A). PS particles of 10µm diameter could not be detected in 50µl plasma 0.5, 2 and 24h after the first administration (Fig. 2A). Furthermore, neither 1µm nor 10µm particles were detected in the cellular fraction of 200µl of blood at 0.5, 2 and 24h after the first administration (data not shown).

**Figure 2.**
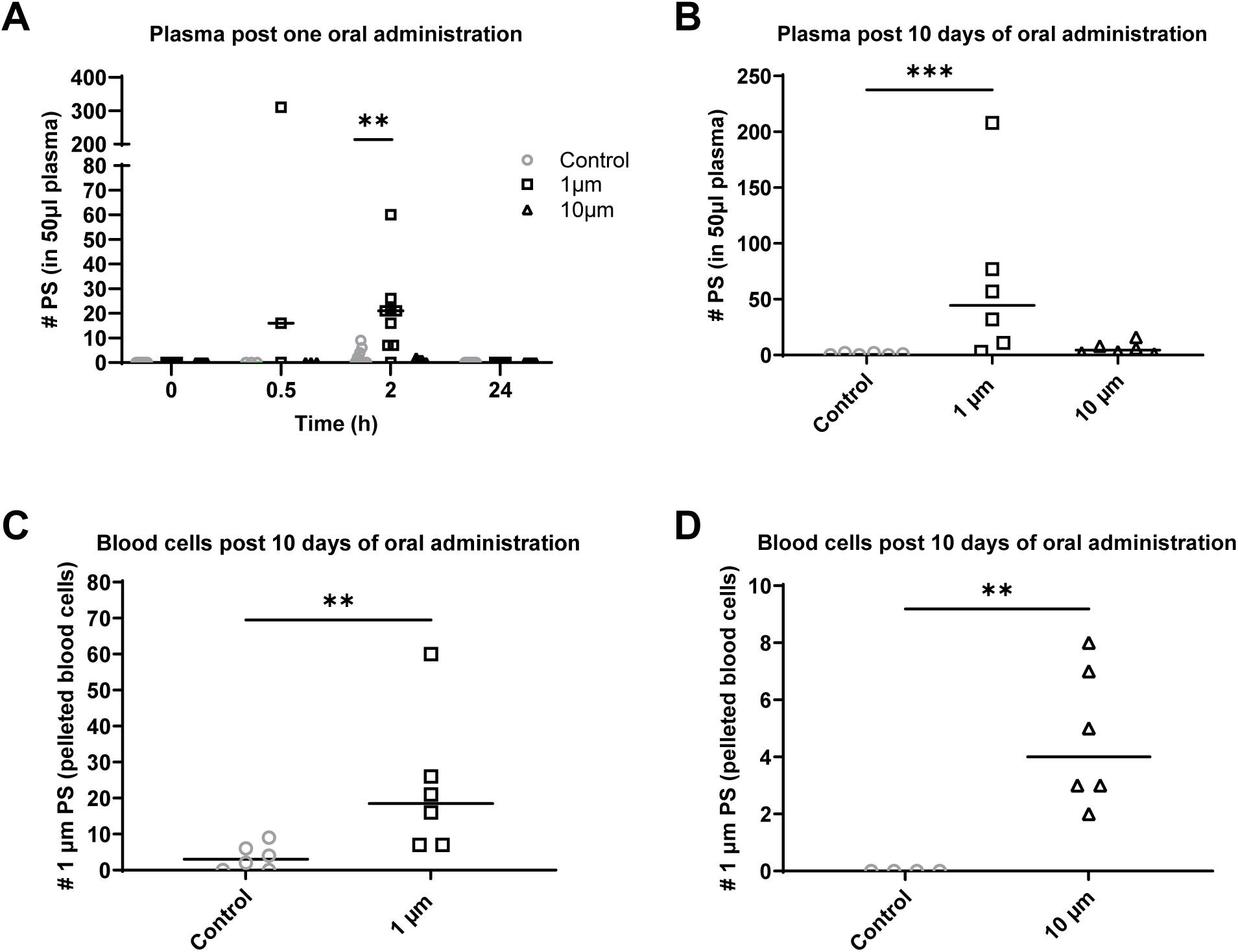
Presence of 1µm and 10µm PS particles in blood after oral administration Number of 1µm and 10 µm PS particles detected in plasma post one oral administration (A) and 10 days post oral administration (B). Number of 1µm PS particles detected in pelleted blood cells 10 days post oral administration (C). Number of 10µm PS particles detected in pelleted blood cells 10 days post oral administration (D).

After 10 days of oral administration, 1µm PS particles could be detected in both plasma and pelleted cells (Fig. 2B and C). The 10µm PS particles were only significantly higher than controls in pelleted cells (Fig. 2D) but not in plasma (Fig. 2B). A median of 45 and 6 particles of 1 and 10 µm diameter respectively were detected in 50 µl of plasma (Fig. 2B). The pelleted cellular fraction contained a median of 19 (Fig. 2C) and 4 (Fig. 2D) particles of 1 and 10µm diameter, respectively.

Taking into account an equal distribution of particles in blood, an average blood volume of 2.4 ml per mouse and 55% plasma on total blood volume^26^, we calculated that on average a mouse at day 10 had 1291 and 180 of 1 and 10µm PS respectively circulating in their blood.

Sections of cryopreserved liver showed few to none 1 and 10µm particles 2 and 24 hours after the first oral administration, whereas both 1 and 10 µm fluorescent PS could be observed after 10 days of oral administration (Fig. 3). To rule out any possibility that the histological sections were contaminated during organ removal, we set up intravital 2-photon microscopy experiments with the aim to determine the presence of particles deep within the liver. Intravital imaging of the liver showed that particles of 1 and 10µm were visible in the liver already after the second oral administration (Video 1 and 2).

**Figure 3.**
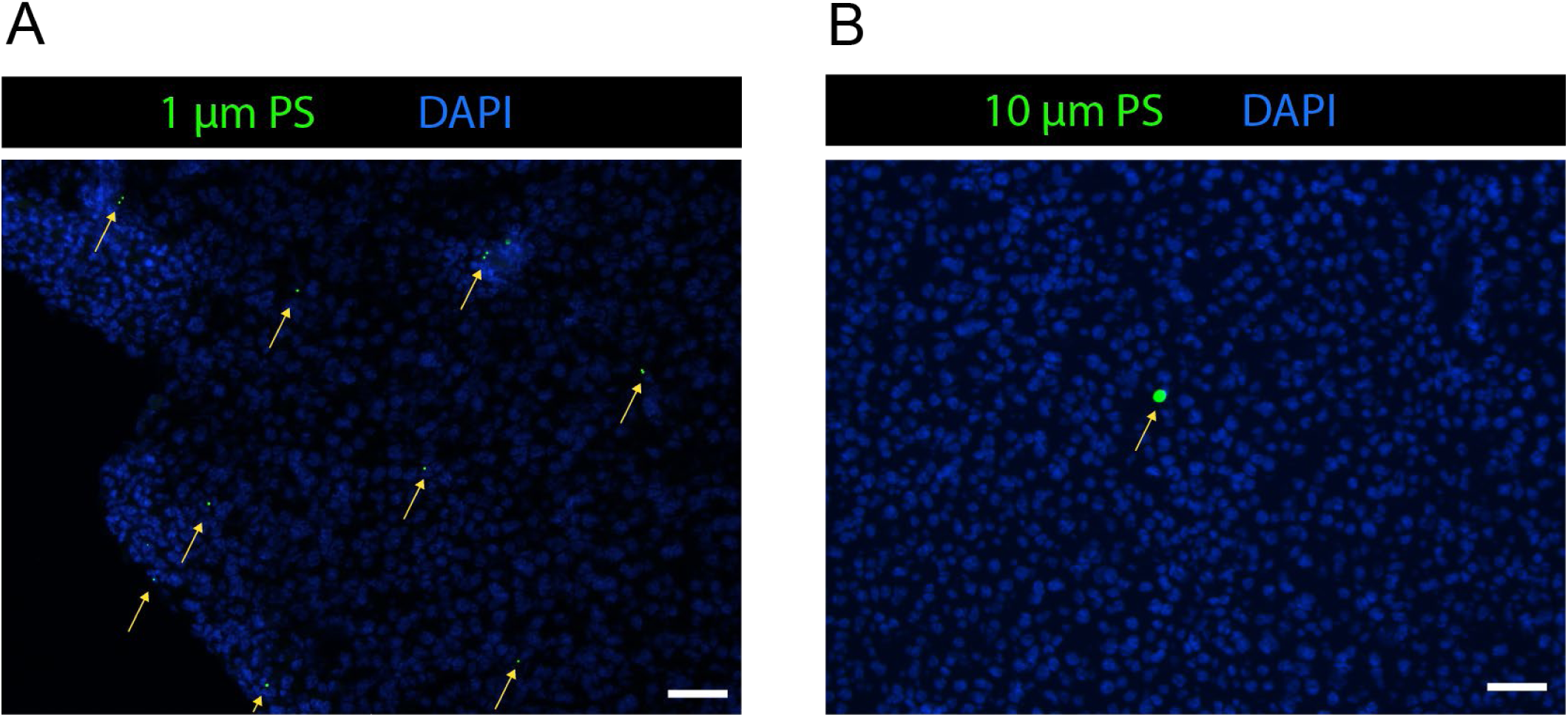
Presence of 1µm and 10µm PS particles in liver after oral administration Liver section showing green 1µm (A) and 10µm (B) PS particles appointed by yellow arrows. Nuclei are stained blue using DAPI. Scale bar is 50 µm.

### Mouse neutrophils phagocytose 1µm PS in vivo after oral and intravenous administration

The evidence collected in our animal studies pointed out that polystyrene particles up to 10µm diameter penetrated the gut barrier and translocated to the blood and liver. When a biological foreign material penetrates the body, this is engulfed by immune cells by an active process called phagocytosis. This process requires the recognition of patterns on the particles by receptors on immune cells. Therefore, we first wanted to establish whether immune cells would at all phagocytose inert PS particles in vitro. In the presence of mouse serum, 1µm PS were readily phagocytosed by mouse neutrophils, while no phagocytosis was observed for 10µm PS (data not shown).

To observe the PS-neutrophil interactions in vivo when the particles reach the liver via the blood circulation, we orally administered either 1 or 10µm pristine fluorescent polystyrene particles for 2 days and prepared the animals for intravital microscopy immediately after the second administration. Neutrophil interactions with 1µm (Video 1) and 10µm (Video 2) PS could be detected although no particle was engulfed or removed from the initial location where they were detected.

Since the number of PS in the liver was limited after oral administration, we additionally performed a proof of principle experiment where 1µm PS were intravenously injected. In this model 1µm PS also reached the liver via the blood circulation, but more events could be recorded. We observed the phagocytosis of 1µm PS particles within the vascular bed of the liver. The neutrophils were observed to catch, engulf and carry the polystyrene particle (Fig. 4 and Video 3).

**Figure 4.**
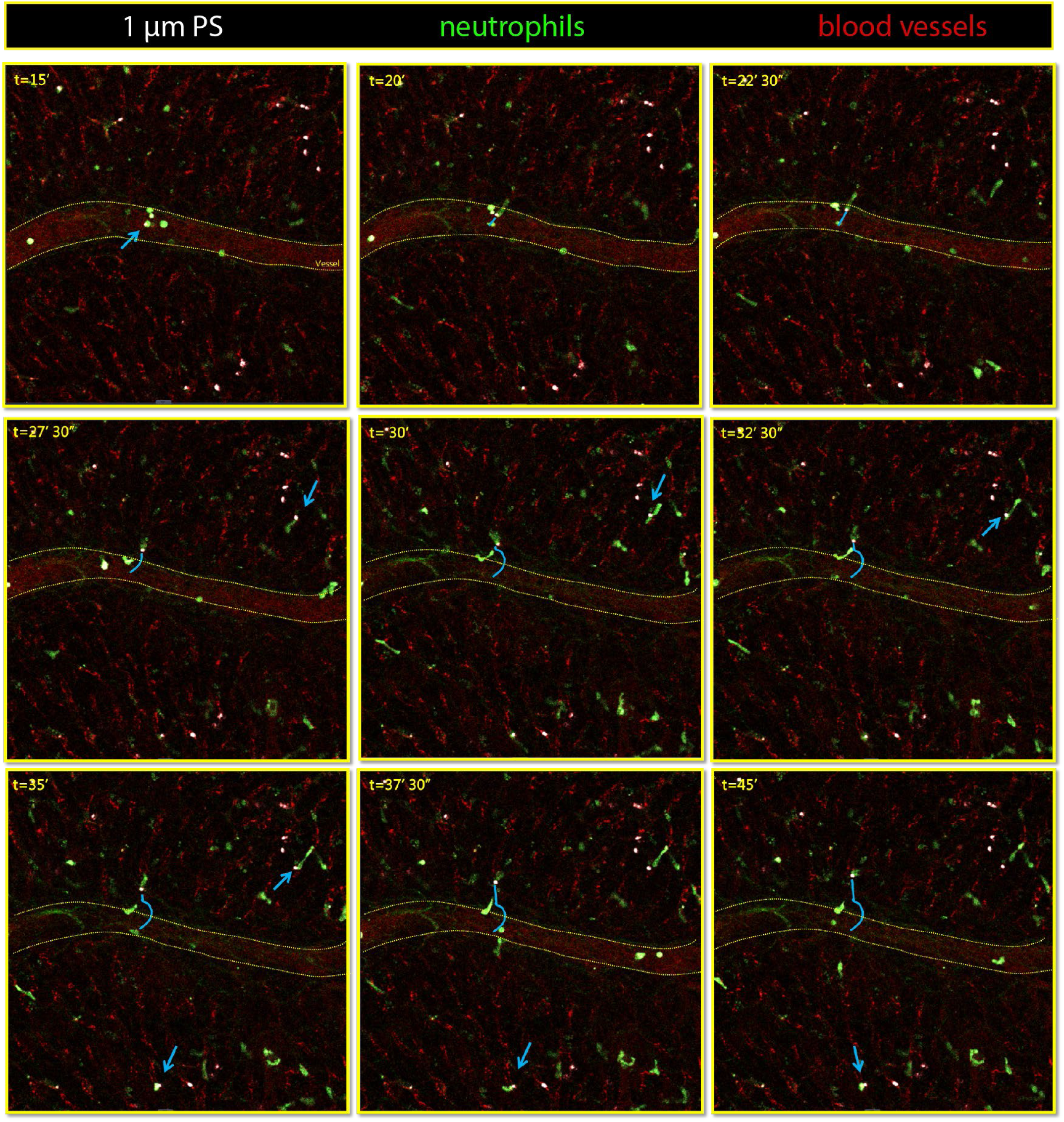
In vivo neutrophil phagocytosis of intravenously injected 1µm PS in livers of mice LysM-GFP mice bearing green neutrophils were intravenously injected with red 1µm PS particles (white in picture) and livers were intravitally imaged. Arrows indicate neutrophils interacting with the 1µm PS particle in vivo. Blue line indicates the track of a 1µm PS particle inside a neutrophil. Scale bar is 50 µm.

Human primary neutrophils and monocytes phagocytose plasma coated polystyrene particles The evidence of PS translocation over the intestinal barrier after oral administration and the phagocytosis of PS in vivo in mice prompted us to investigate potential interactions between professional phagocytes and microplastics in human relevant models in vitro. We first determined the minimal requirements for human primary neutrophils to phagocytose 1 and 10µm PS particles in 2D in vitro cultures.

Phagocytosis of 1 and 10µm PS particles was monitored by live imaging of neutrophils on a confocal microscope. To observe phagocytosis in freshly isolated neutrophils, f-MLF-mediated chemokinesis was necessary to induce contact with PS as neutrophils were otherwise immobile (data not shown). Pre-incubation of both 1 and 10µm PS with plasma is also indispensable for phagocytosis in 2D (Fig. 5A and Video 4-6). Coating with human serum albumin, which accounts for approximately 55% of proteins found in plasma, was not sufficient to induce phagocytosis (data not shown). In addition to confocal microscopy we also quantified phagocytosis in time using an Incucyte microscope in an incubator by measuring the overlapping area of green fluorescent PS and red stained phagocytes (Fig. 5B) further confirming plasma coating is necessary for phagocytosis to occur. Monocytes did not need any further stimulus and their spontaneous motility was enough to induce contact between cells and PS particles promoting phagocytosis in 2D. Pre-incubation of both 1 and 10µm PS with plasma was, however, also indispensable for phagocytosis by monocytes (data not shown).

**Figure 5.**
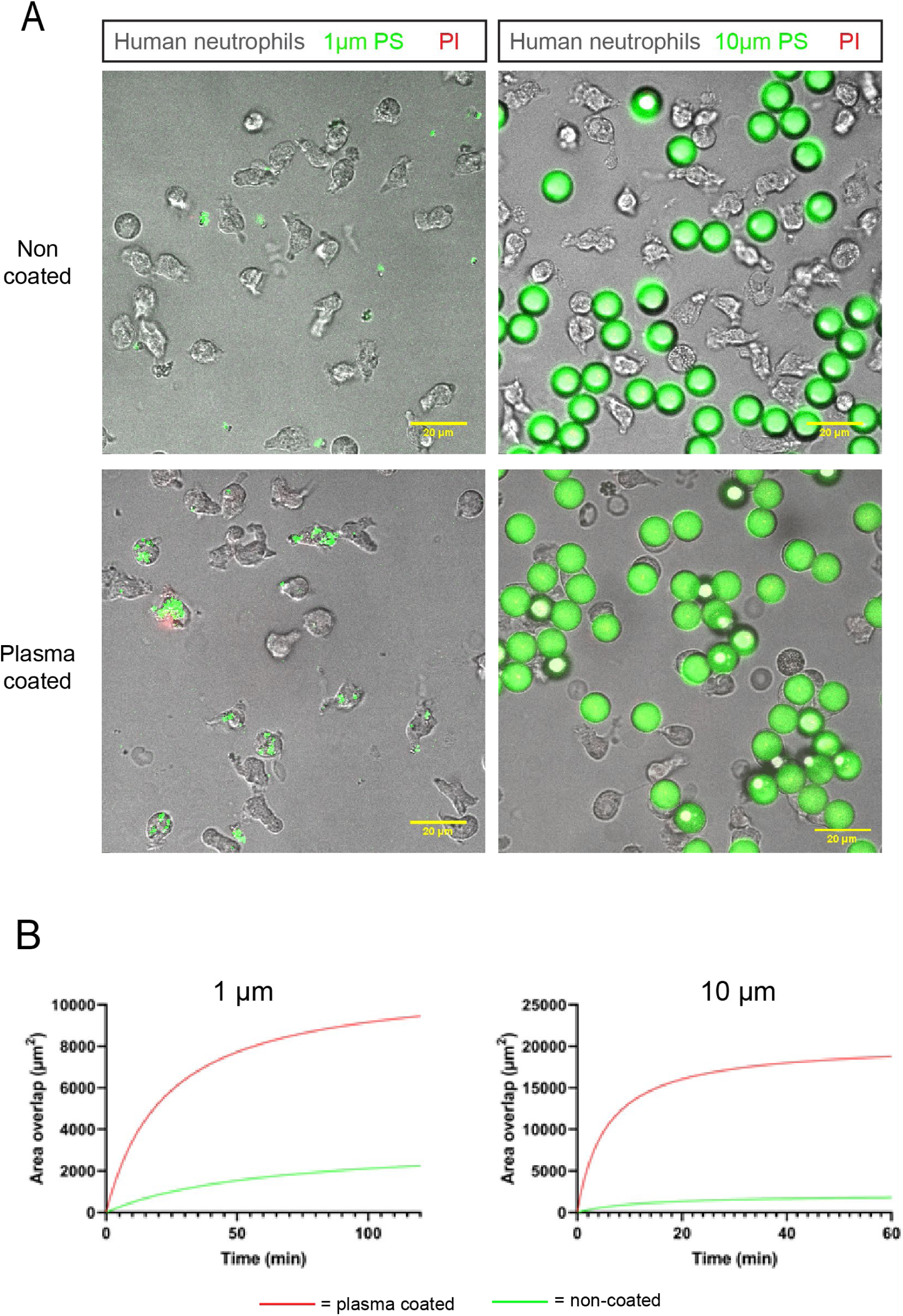
Phagocytosis of uncoated and coated 1µm and 10µm PS particles by human neutrophils (A) Phagocytosis of 1 and 10µm PS particles (green) was monitored by live imaging of f-MLF stimulated neutrophils (grey) on a confocal microscope. PS particles were either uncoated or pre-incubated with plasma. Scale bar is 20 µm. (B) Phagocytosis of uncoated (green line) and plasma-coated (red line) of 1 and 10µm PS particles quantified in time using an Incucyte microscope by measuring the overlapping area of green fluorescent 1 and 10µm PS with red stained neutrophils.

In 2D in vitro cultures, PS particles gradually settle down onto the layer of phagocytes at the bottom of the well. Therefore, we also performed a phagocytosis assay in 3D collagen gels to better mimic the in vivo situation. Phagocytosis of 1 and 10µm PS by neutrophils was also observed in 3D collagen gels. Plasma coated particles were promptly phagocytosed by neutrophils whereas particles which were incubated with only incubation buffer were mostly ignored (Fig. 6C).

**Figure 6.**
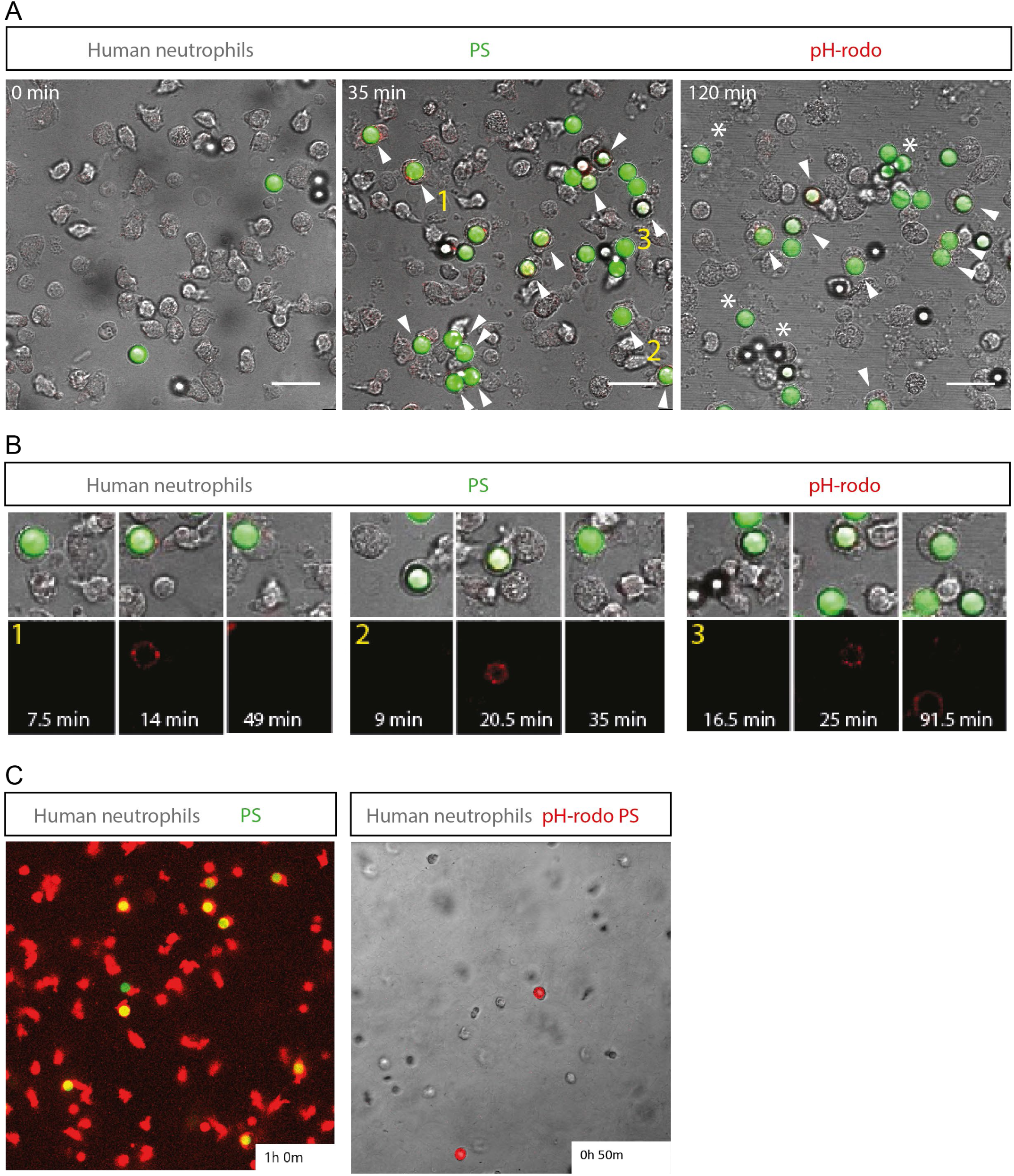
Confirmation of phagocytosis of 1µm and 10µm PS particles in 2D and 3D by pH-rodo (A) Fluorescent 7.3 µm beads were labelled with pH-rodo and added to isolated human neutrophils stimulated with f-MLF. Timelapse confocal microscopy images are shown. Microplastics phagocytosed by neutrophils are shown by arrows and * depicts neutrophil death. Scale bar is 20 µm. (B) The signal of pH-rodo is shown in detail for 3 cells marked 1,2,3 in A. Images right after phagocytosis, at the first timepoint of acidification and at the moment acidification disappears are shown. (C) Left panel depicts phagocytosis of plasma-coated 10µm PS (green) by fMLF-stimulated neutrophils (red) in a maximum projection of a 3D collagen gel after 1 hour. Right panel shows pH-rodo staining of plasma-coated 10µm PS (red) internalized by fMLF-stimulated neutrophils (green) in a maximum projection of a 3D collagen gel at 50 minutes.

To confirm PS internalization, we made use of pH-rodo-coated PS. Upon phagocytosis, the particle resides in a membrane coated vesicle inside the cellular cytoplasm called a phagosome. This vesicle fuses with lysosomes to form phagolysosomes, which causes a decrease in pH within the phagolysosome as a mechanism to stop bacterial growth. pH-rodo-coated PS were additionally pre-incubated with pooled serum and incubated with primary human neutrophils. A decrease in pH, associated with maturation of the phagolysosome, was observed after internalization of the PS both in 2D (Fig. 6A and B) and 3D (Fig. 6C).

### Phagocytosed polystyrene leads to early neutrophil and monocyte cell death

We exposed human primary neutrophils (Fig.7) and monocytes (Fig.8) to different weights of plasma-coated 1, 5 and 10µm PS particles and measured their cell death over time by fluorescence microscopy. Obviously, the same weight of 1µm PS particles represents a 1000 times higher particle number than that same weight of 10µm PS particles. Therefore, we also determined the number of PS per cell (PS:cell ratio) for each weight (Fig.7B and 8B).

**Figure 7.**
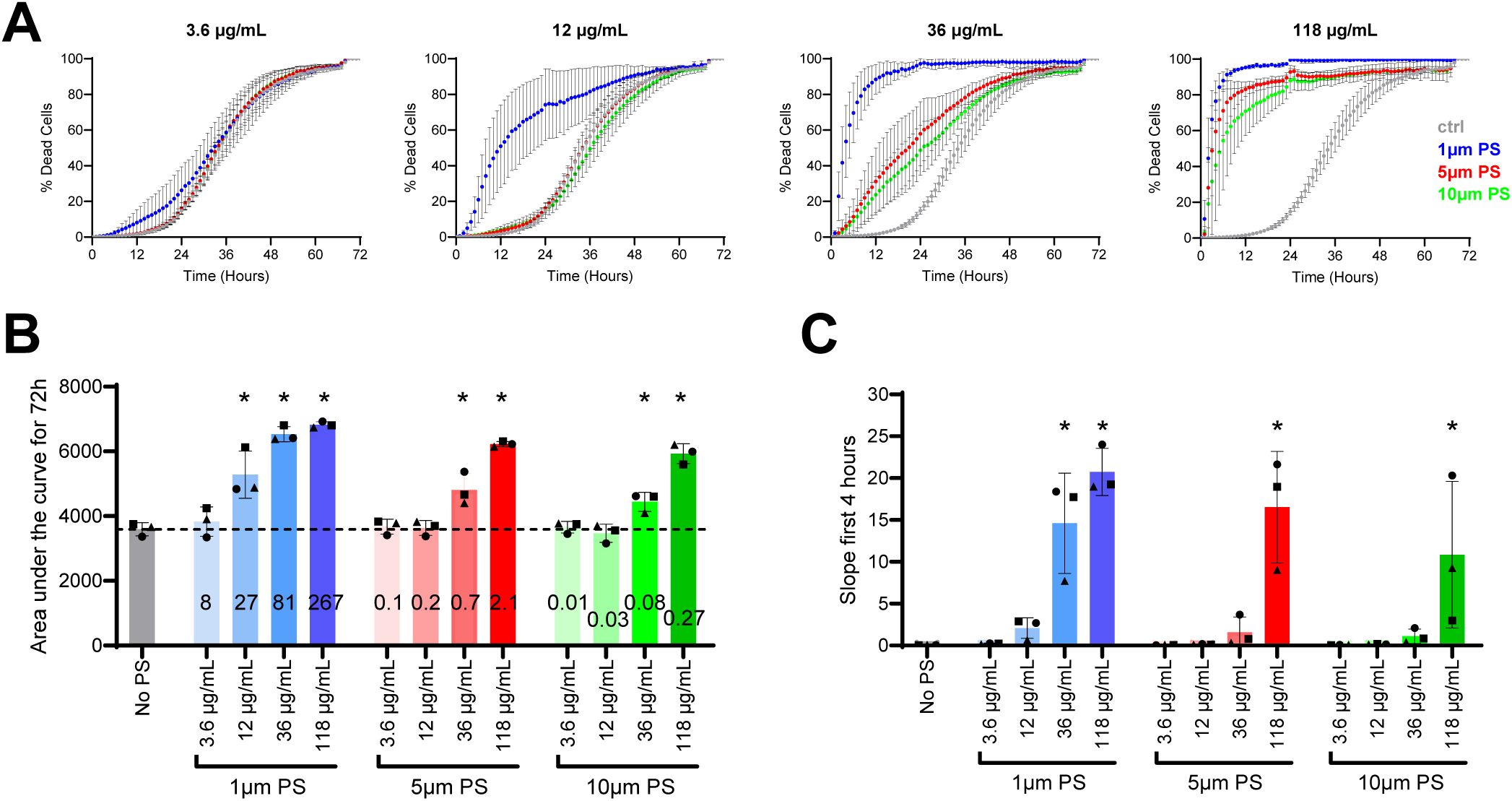
Kinetic neutrophil cell death assay in response to different concentrations of 1µm, 5µm and 10µm PS particles (A) Timelapse microscopy on the Incucyte detected the MFI of PI as a measure for dead neutrophils in response to different concentrations of 1µm, 5µm and 10µm PS particles and controls. Results shown are means and standard deviations of 3 biological replicates. (B) Area under the curve of graphs depicted in A for individual biological replicates. Numbers in the bars are the ratio of PS:cell for that specific weight concentration. Dotted line is the mean of controls. (C) Slope of the first four hours of graphs depicted in A for individual biological replicates. Significance compared to control is depicted with an asterisk.

**Figure 8.**
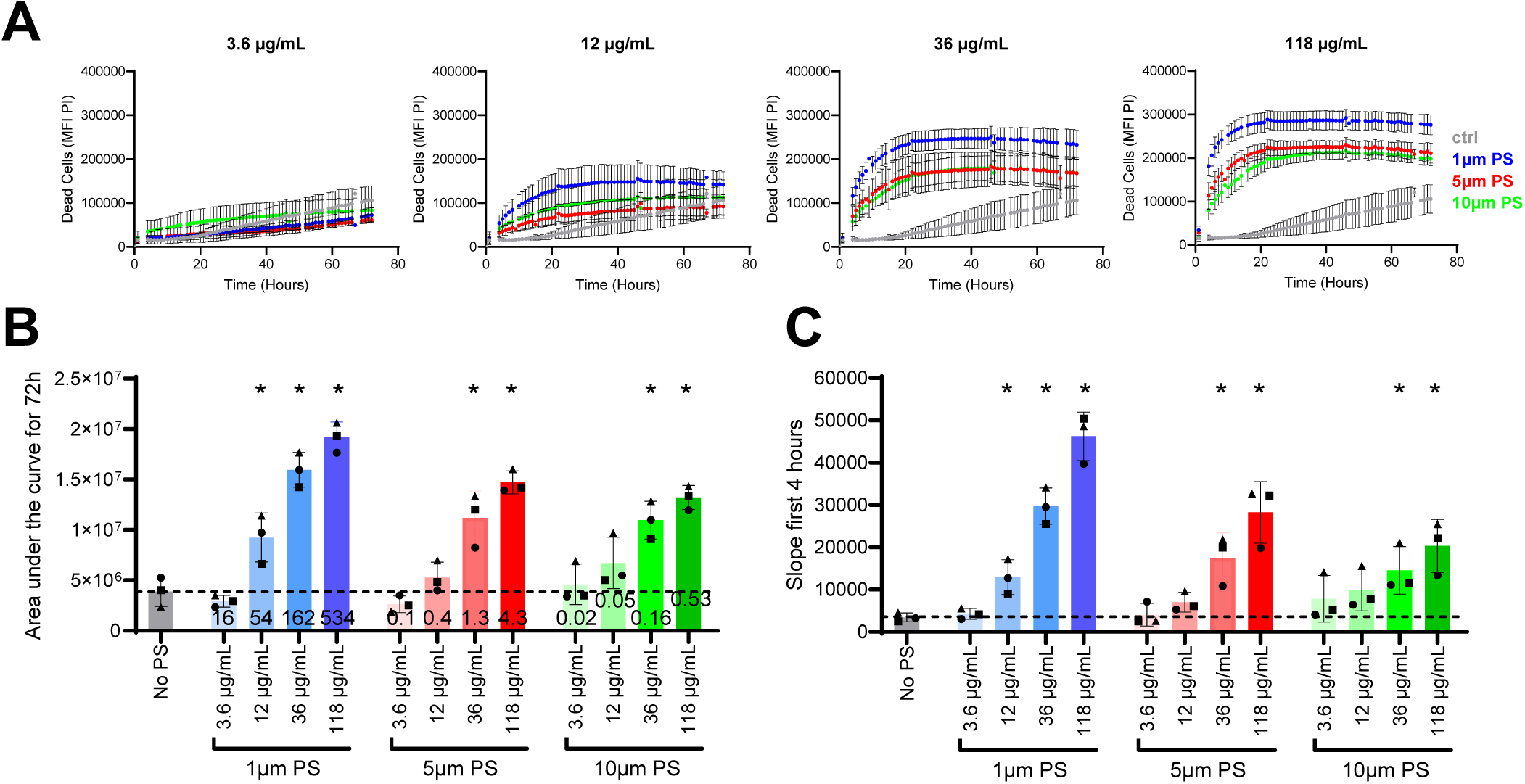
Kinetic monocyte cell death assay in response to different concentrations of 1µm, 5µm and 10µm PS particles (A) Timelapse microscopy on the Incucyte detected the MFI of PI as a measure for dead monocytes in response to different concentrations of 1µm, 5µm and 10µm PS particles and controls. (B) Area under the curve of graphs depicted in A for individual biological replicates. Numbers in the bars are the ratio of PS:cell for that specific weight concentration. Dotted line is the mean of controls. (C) Slope of the first four hours of graphs depicted in A for individual biological replicates. Results shown are means and standard deviations of 3 biological replicates. Significance compared to control is depicted with an asterisk.

Plasma-coated 1µm PS did not induce any significant cell death when added at a particle:cell ratio of 8:1 but did induce significant cell death when added at a ratio of 27:1 or higher (Fig.7). Plasma-coated 5µm PS already induced significant cell death when added in a ratio of 0.7:1 or higher and plasma-coated 10µm in a ratio of 0.08:1 (Fig.7). Cell death is therefore not solely weight based but more importantly number and size based. (Fig.7). An early significant increase of neutrophil and monocyte cell death could, however, be detected for all three sizes (Fig.7 and 8).

### Phagocytosis of PS is essential to induce early neutrophil death

The increased cell death in neutrophils and monocytes treated with plasma-coated 1, 5 and 10µm PS prompted further investigation to determine if phagocytosis of PS was necessary to trigger cell-death. Interestingly, cells exposed to non-coated or HSA-coated PS, which we showed earlier were not phagocytosed (Fig. 6), did not show any significant increase in cell death compared to controls not receiving PS (Fig. 9). In addition, we exposed isolated neutrophils to plasma-coated 10µm PS and incubated them for 1h in presence of f-MLF to allow for engulfment of these particles. After 1 hour, cells were prepared for flow cytometry cell sorting and three populations of events were identified: neutrophils containing a 10µm fluorescent PS, non-fluorescent neutrophils and fluorescent 10µm PS which were not associated with cells. Fluorescent 10µm PS containing neutrophils and non-fluorescent control neutrophils were seeded in different wells. In the well containing non-fluorescent cells the same number of fluorescent PS was added at a later step. At this time, neutrophils were not motile since f-MLF was not replaced in the wells and no further phagocytosis was observed (Fig. 9). An early increase in cell death was observed only in neutrophils that phagocytosed the 10µm PS before cell sorting, while non-fluorescent neutrophils co-incubated with a comparable number of PS did not die earlier than controls (Fig. 9). Together these data indicate that phagocytosis of 10µm PS is essential to induce early cell death. Furthermore, this experiment demonstrated the effect is particle related and ruled out that chemicals leaching from the particles caused the effect.

**Figure 9.**
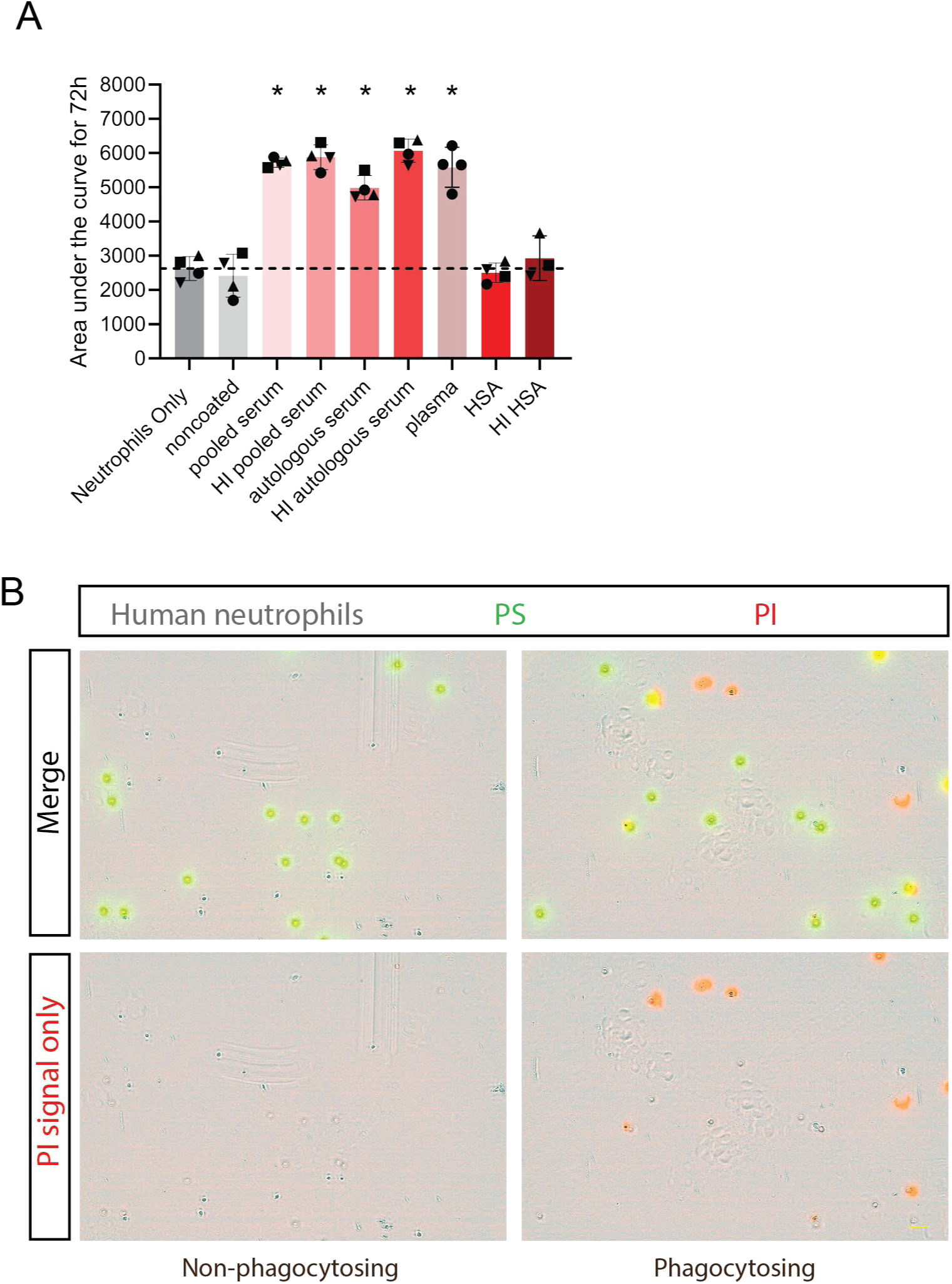
Phagocytosis of PS particles is essential for the induction of neutrophil cell death (A) Timelapse microscopy on the Incucyte detected the MFI of PI as a measure for dead neutrophils in response to differentially coated 10µm PS particles in a PS:cell ratio of 1. The AUC of this kinetic curve was determined and compared to controls without PS . Results shown are means and standard deviations of 4 biological replicates. Dotted line is the mean of controls. Significance compared to control is depicted with an asterisk. (B) fMLF stimulated neutrophils (grey) that did or did not phagocytose 10µm PS particles (green) were sorted by flow cytometry. Neutrophils which did not phagocytose 10µm PS particles were supplemented with equal numbers of extracellular 10µm PS particles which were not further phagocytosed due to further the absence of fMLF.

**Figure 10.**
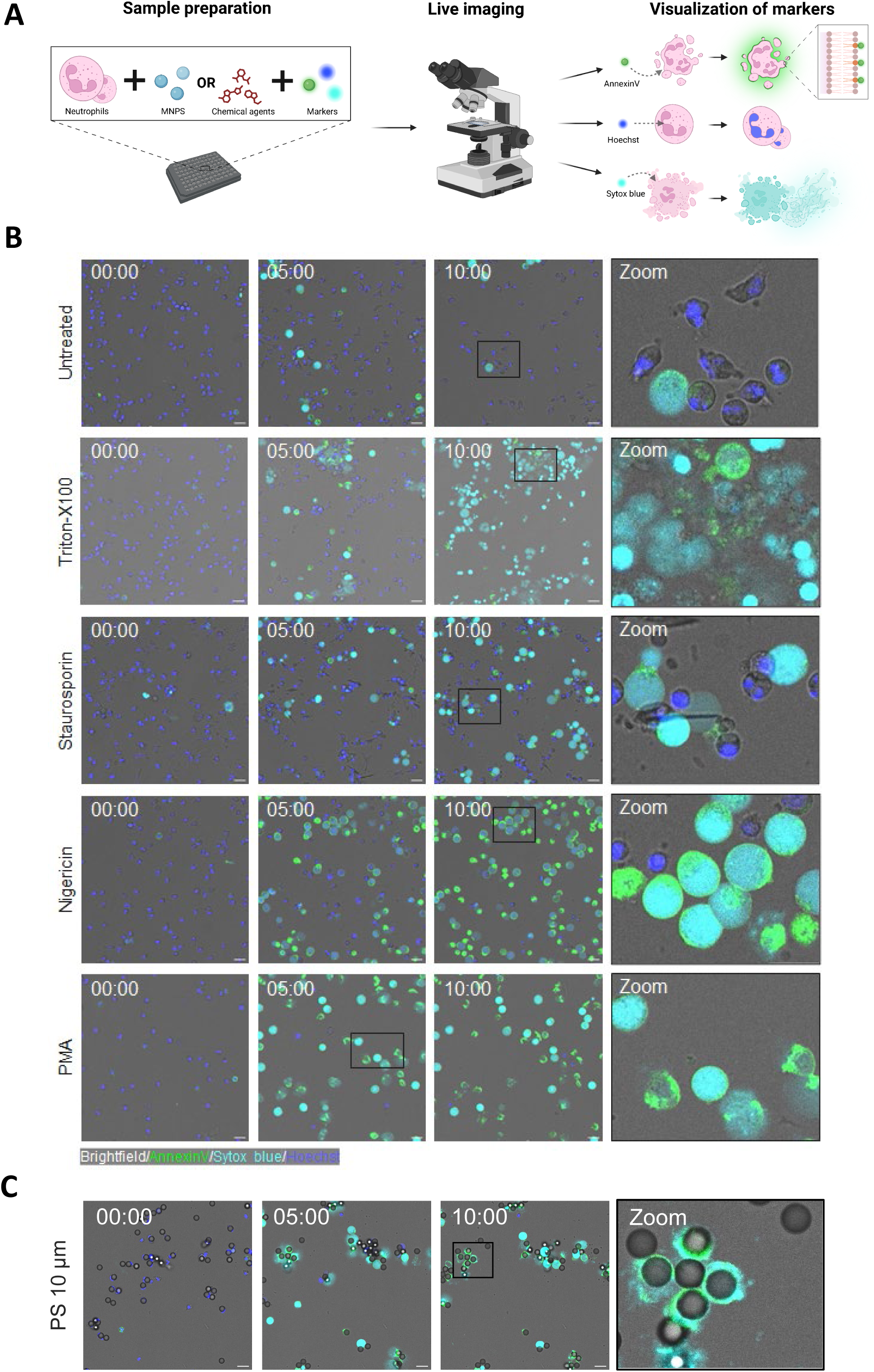
Phagocytosis of PS particles is essential for the induction of neutrophil cell death Isolated neutrophils from healthy donors were either untreated or exposed to various chemical agents in the presence of AnnexinV (green), Sytox blue (cyan), and Hoechst (blue). The neutrophils were live imaged every 10 minutes for over 10 hours to visualize cell death over time. (A) Schematic representation of the performed cell death assay, created with BioRender. (B) Representative images of real-time cell death of neutrophils when either untreated or exposed to 0.05% (w/v) Triton-X100, 10 µM Staurosporin, 10 µM Nigericin, or 25 ng/ml PMA at indicated timepoints in hours. (C) Neutrophil cell death after exposure to 137.44 μg/ml 10 µm PS (1:1 neutrophil to PS ratio). For all the conditions, the last image is a zoomed in image of timepoint 10h, except for PMA which is zoomed in at the 5h timepoint. The images are representative of 3 independent experiments with 3 different donors. Scale bar 20 µm.

### 10µm PS induce pro-inflammatory neutrophil cell death characterized by extracellular DNA

To visualize and to distinguish the various forms of cell death over time, neutrophils were exposed to chemical agents or coated 10µm PS in the presence of fluorescent cell death markers, such as AnnexinV (binds extracellularly exposed phosphatidylserine of early apoptotic cells and intracellular phosphatidylserine in permeable dead cells), Hoechst (cell-permeable dye which binds DNA to stain the nucleus of live cells), and Sytox blue (binds DNA of dying, permeable cells) (Fig.10A).

Neutrophils were exposed to various chemical agents to induce different forms of cell death as a control for the cell death induced by 10µm PS (Fig.10B). Few neutrophils died in the untreated controls allowing the influx of Sytox blue into the cell where most of the DNA remained within the cell, which implies non-inflammatory cell death. Triton-X100 was used as a necrosis inducer, which was characterized by the disintegration of the cell membrane and an increase in Sytox blue signal. Within the imaged time, some of the DNA was still present within the nucleus and some DNA was extracellularly present after exposure to Triton-X100. The neutrophils treated with the apoptosis inducer Staurosporin formed blebs and needle-like structures over time. Most dying cells from Staurosporin demonstrate Sytox blue within the cell that is surrounded by an AnnexinV+ cell membrane, indicating non-inflammatory cell death. Both Nigericin and PMA are known to induce extracellular DNA either via NETosis or pyroptosis^27^. However, upon treatment with nigericin, neutrophils had a more swollen appearance and most of the DNA remained intracellular. In contrast, neutrophils exposed to PMA demonstrated consistently more extracellular DNA.

When neutrophils were treated with 10μm PS particles, the particles were phagocytosed upon which swelling of the cell was noted and eventually extracellular DNA was released (Fig.10C and Video 7). The form of cell death is comparable with the cell death seen in PMA-treated neutrophils, both leading to an increase in extracellular DNA based on the Sytox blue staining, indicating pro-inflammatory cell death. After cell death, the AnnexinV+ cell membrane remains attached to the 10 μm PS particles. This data indicates that neutrophil exposure to 10 μm PS particles results in increased pro-inflammatory cell death.

## Discussion

An increase in the spreading of plastic pollution worldwide and recent evidence showing that plastic particles can be detected in the blood and brain of human beings^28,29^, urged the scientific community to rapidly define the toxicological hazard and the risk for these particles in humans^30^. The immune system, and especially the innate branch, might likely be the first to encounter these microparticles when introduced into our body via the respiratory and the gastrointestinal tracts. In particular, specialized phagocytic cells such as neutrophils and macrophages might be the first cells to interact with plastic microparticles as shown already for other particle matter such as silica^31–34^.

In this study the first aim was to confirm the translocation of 1 and 10µm PS over the intestinal barrier. Hence, we orally administered PS of 1 and 10µm diameter to mice for 10 days and demonstrated that both particles crossed the epithelial gut barrier and were found circulating in the blood of these mice. The 1µm PS were already detectable in plasma of some mice 30 minutes after the first oral administration whereas 10µm PS were never detected in plasma but only detected in pelleted cells after ten days of PS administration. In this study, we orally administered the same weight and thus numerically 1000 times more of 1µm compared to 10µm PS particles to mice. We did, however, not detect 1000 times more 1µm PS particles in pelleted blood cells or livers after ten days of PS administration when compared to 10µm PS. This suggests that 10µm PS particles surprisingly might pass the intestinal barrier relatively more efficient than 1µm PS.

After two days of oral administration, 1 and 10µm PS were detected deep in the liver using intravital microscopy of mice under anesthesia. Furthermore, in vivo phagocytosis of 1µm but not 10µm PS was recorded in the liver of mice that were intravenously injected with PS particles. These data further confirm PS particles reach the circulation after oral administration in mice and establish phagocytosis of PS particles occurs in vivo. It is noteworthy that mouse experiments might produce results which cannot be extrapolated to humans, as we showed that mouse neutrophils did not phagocytose PS microplastics of 10µm, while human neutrophils can readily phagocytose microplastics of this size.

The experiments conducted in vitro with the use of human neutrophils and monocytes showed that plasma coating greatly increased the amount of microplastics that were phagocytosed. This suggests that some form of opsonisation occurs during the incubation with plasma, potentially by complement and/or antibodies. Furthermore, neutrophils required fMLF mobilization while monocytes did not. It is of interest that pre-coating or priming was not necessary to observe phagocytosis by neutrophils in vivo in mice, suggesting that the coating of microplastics also occurs in vivo. Importantly, environmental microplastics are surrounded by a “corona” of non- and biological matters with potential biological effects on cells^35,36^. The environmental corona represents a dense concentrate of potential toxicants, immunostimulants as well as a potential source of new antigens^37–39^. Efforts should be spent to clarify how the eco-corona is affected when entering a biological system via different routes.

Importantly, neutrophil and monocyte cell death occurred a few hours after phagocytosing a single coated 10µm PS while PS of 1µm needed to be administered at a ratio of 27 particles per neutrophil or 54 particles per monocyte to induce significant neutrophil cell death. The similarities between the effects observed after exposure to PS in 2D and 3D confirms that this behaviour was not caused by neutrophils being on an artificial 2D plane and further justifies the extrapolation that microplastics might be phagocytosed by neutrophils in humans in vivo. Furthermore, the 3D model is a valuable tool to measure the effects of buoyant microplastics in future experiments.

Our experiments incontrovertibly demonstrated that the mere incubation of PS with neutrophils was not sufficient to induce cell death. When preventing phagocytosis by not coating the PS particles or by not inducing cell motility by fMLF, cell death did not occur. This observation rules out a role for chemical leachates in the observed toxicity for PS. The experiments underlined an existing threshold in weight uptake beyond which neutrophils and monocytes induced cell death. Neutrophils needed to be exposed to a 1µm PS:cell ratio of 27 to induce toxicity, which is unlikely to occur in vivo. On the other hand, for 10µm PS only one particle per cell was already enough to induce neutrophil cell death. It is noteworthy that the current estimates of microplastics in blood are determined based on their weight, but their size range is unknown^19^.

Experiments in our study were done with commercially available perfectly spherical PS particles that are being produced ‘bottom-up’, while environmental microplastics will consist of broken-down plastics of multiple materials, shapes and sizes. Future experiments should be aimed at testing neutrophil and monocyte cell death after exposure to more environmentally relevant particles.

In conclusion, this study underlined how PS ingestion led to the translocation and systemic circulation of these particles through the blood and the liver. Phagocytosis of microplastics by immune cells occurred in vivo in mice as well as in human phagocytes in vitro, leading to inflammatory cell death in human neutrophils and monocytes. Future studies should compare different plastic materials, shapes and sizes as well as different eco-corona’s.

Since humans come into contact with microplastics in their day-to-day lives and they are omnipresent within the environment, it is of utmost importance to fully elucidate the effects of microplastics on the health of both humans and animals. Our results hint at the possibility that microplastics can have adverse effects on the immune system and on human health. It is, however, difficult to say how this observed cell death would translate to the level of a tissue or a human being as a whole. The increased death of neutrophils upon microplastic exposure can possibly lead to pro-inflammatory reactions caused by the extracellular DNA and other DAMPS released from dead neutrophils. This might recruit other immune cells which start phagocytosing the released microplastic microplastics. Alternatively, the released microplastics might be left behind inside the tissue where the phagocytes die. Because PS is considered a chemically and enzymatically inert material^40^, the permanence of these particles in biological systems may trigger biological responses. More research is needed to further elucidate the potential effect of the accumulation of microplastics in the human body.

## Acknowledgements

This research received funding from the EC Horizon 2020-project POLYRISK [Grant ID 964766], the ZonMw/Health Holland project MOMENTUM [Grant ID 458001101] and the Netherlands Organisation for Scientific Research (NWO) in the context of a private public partnership Technology Area COAST [Grant ID 053.21.112].

---

Video 1. Intravital microscopy of neutrophil (green) interactions with orally administered 1µm PS (white) in livers of mice

Video 2. Intravital microscopy of neutrophil (green) interactions with orally administered 10µm PS (white) in livers of mice

Video 3. Intravital microscopy of neutrophil (green) phagocytosis of intravenously injected 1µm PS (white) in livers of mice

Video 4. Live confocal microscopy imaging of human neutrophils (grey), PI to stain DNA of dead cells (red) and non-coated 10µm PS (green)

video 5. Live confocal microscopy imaging of human neutrophils (grey), PI to stain DNA of dead cells (red) and plasma-coated 10µm PS (green)

video 6. Live confocal microscopy imaging of human neutrophils (grey), PI to stain DNA of dead cells (red) and plasma-coated 1µm PS (green)

Video 7. Live confocal microscopy imaging of human neutrophils (light grey), 10µm PS (dark grey), AnnexinV (green), Hoechst (blue, DNA in living cells) and Sytox blue (cyan, extracellular DNA)

