## Supplementary figures and images for "Microplastics cross the murine intestine and induce inflammatory cell death after phagocytosis by human monocytes and neutrophils"

### Video 1.gif

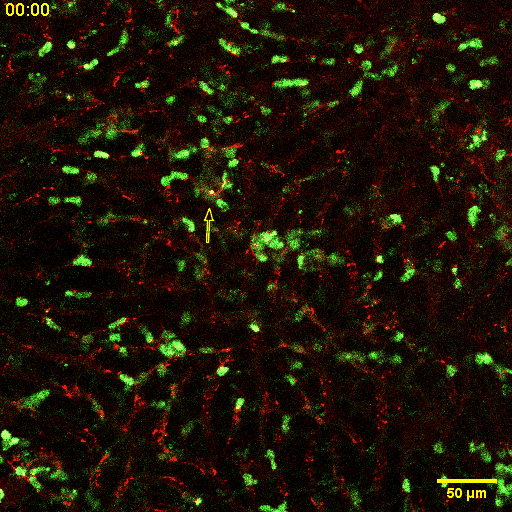

### Video 2.gif

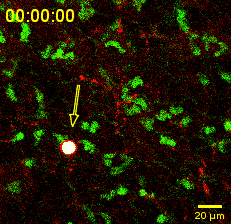

### Video 3.gif

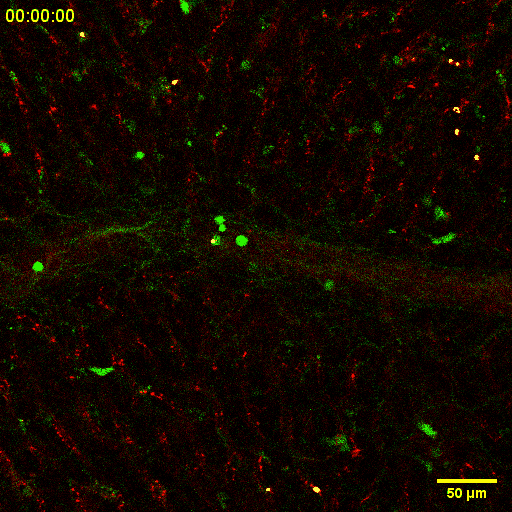

### Video 4.gif

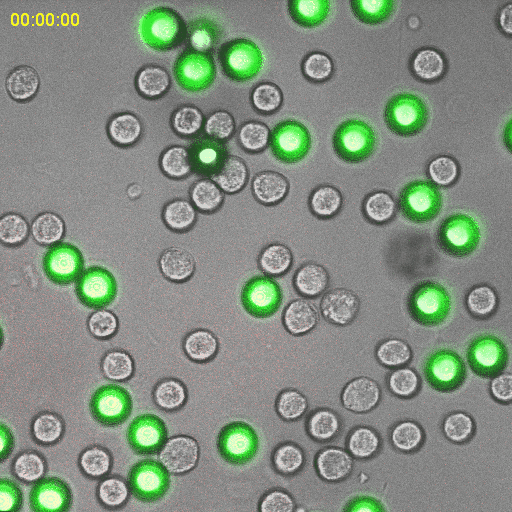

### Video 5.gif

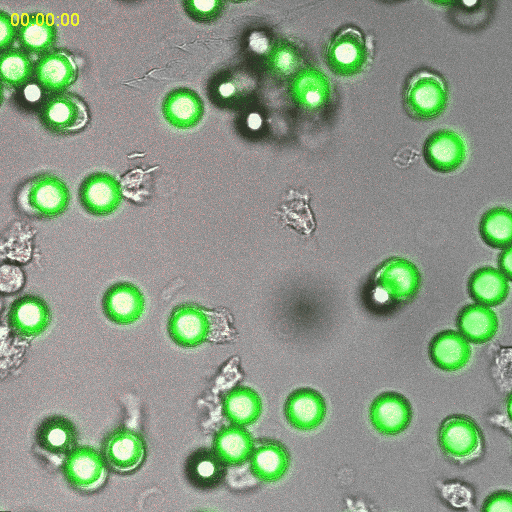

### Video 6.gif

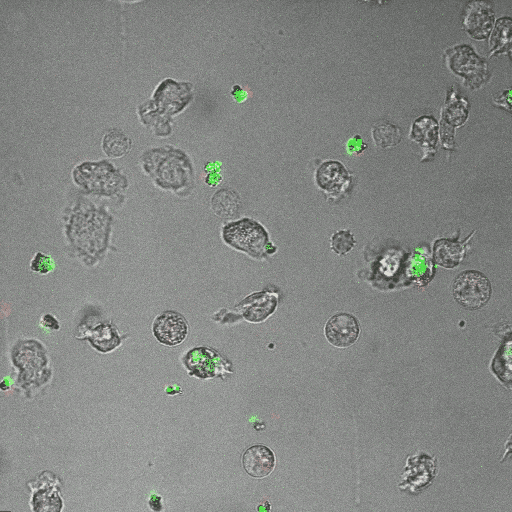

### Video 7.gif

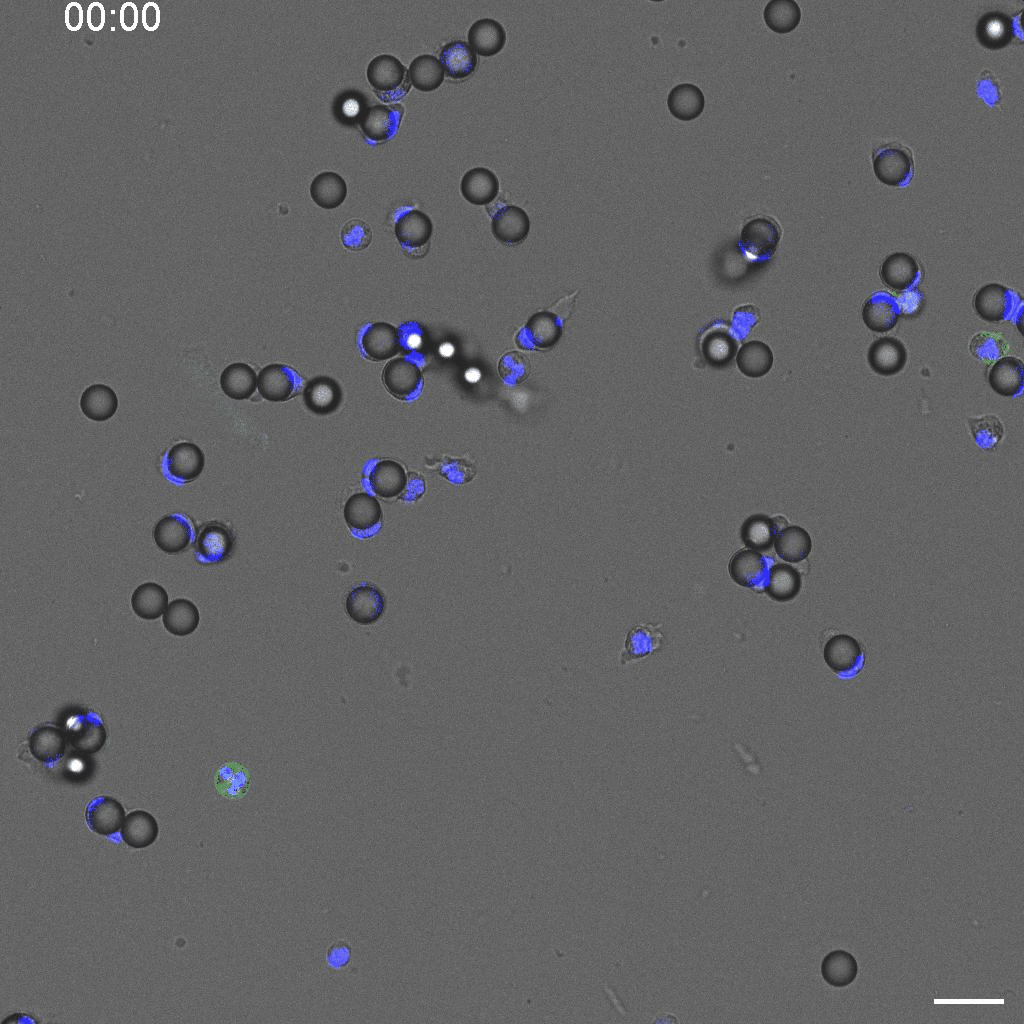
